# Dopamine Neurons Evaluate Natural Fluctuations in Performance Quality

**DOI:** 10.1101/2021.06.23.449657

**Authors:** Alison Duffy, Kenneth W. Latimer, Jesse H. Goldberg, Adrienne L. Fairhall, Vikram Gadagkar

**Author notes:** Correspondence (A.L.F), (V.G.).

## Abstract

Many motor skills are learned by comparing ongoing behavior to internal performance benchmarks. Dopamine neurons encode performance error in behavioral paradigms where error is externally induced, but it remains unknown if dopamine also signals the quality of natural performance fluctuations. Here we recorded dopamine neurons in singing birds and examined how spontaneous dopamine spiking activity correlated with natural fluctuations in ongoing song. Antidromically identified basal ganglia-projecting dopamine neurons correlated with recent, and not future, song variations, consistent with a role in evaluation, not production. Furthermore, dopamine spiking was suppressed following the production of outlying vocal variations, consistent with a role for active song maintenance. These data show for the first time that spontaneous dopamine spiking can evaluate natural behavioral fluctuations unperturbed by experimental events such as cues or rewards.

## INTRODUCTION

Dopamine (DA) is associated with fluctuations in future movements as well as the outcomes of past ones. During spontaneous behavior, DA activity can be phasically activated before a movement (da Silva et al., 2018; Hamilos et al., 2020), or can ramp as an animal approaches reward (Hamid et al., 2016; Howe et al., 2013). DA neurons can also signal a reward prediction error (RPE) during reward seeking, where phasic signals represent the value of a current outcome relative to previous outcomes (Schultz et al., 1997). It remains poorly understood how spontaneous DA activity relates to natural fluctuations in behavior that are independent of experimentally induced rewards or perturbations.

Zebra finches provide a tractable model to study the role of DA in natural behavior. First, they sing with a significant amount of trial-to-trial variability, but the overall stereotypy of the song allows renditions to be accurately compared. Second, they have a discrete neural circuit (the song system) that includes a DA-basal ganglia (BG) loop (Figures 1A and 1B) that is necessary for song learning and maintenance (Brainard and Doupe, 2000; Hisey et al., 2018; Hoffmann et al., 2016; Xiao et al., 2018). Third, BG projecting DA neurons signal performance prediction error (PPE) during singing: they exhibit pauses following worse-than-predicted outcomes caused by distorted auditory feedback (DAF), and they exhibit phasic bursts following better-than-predicted outcomes when predicted distortions do not occur (Figures 1A-1D) (Gadagkar et al., 2016). Yet one limitation of this study was that song quality was controlled with an external sound (DAF, Figure 1C), so it remains unclear if the DA system is simply using song timing to build expectations about an external event (DAF), or if it also evaluates the quality of natural fluctuations (Figure 1E), which would be necessary for natural song learning. Furthermore, this experimental paradigm did not test if DA activity was associated with upcoming syllables, consistent with a premotor signal.

**Figure 1.**
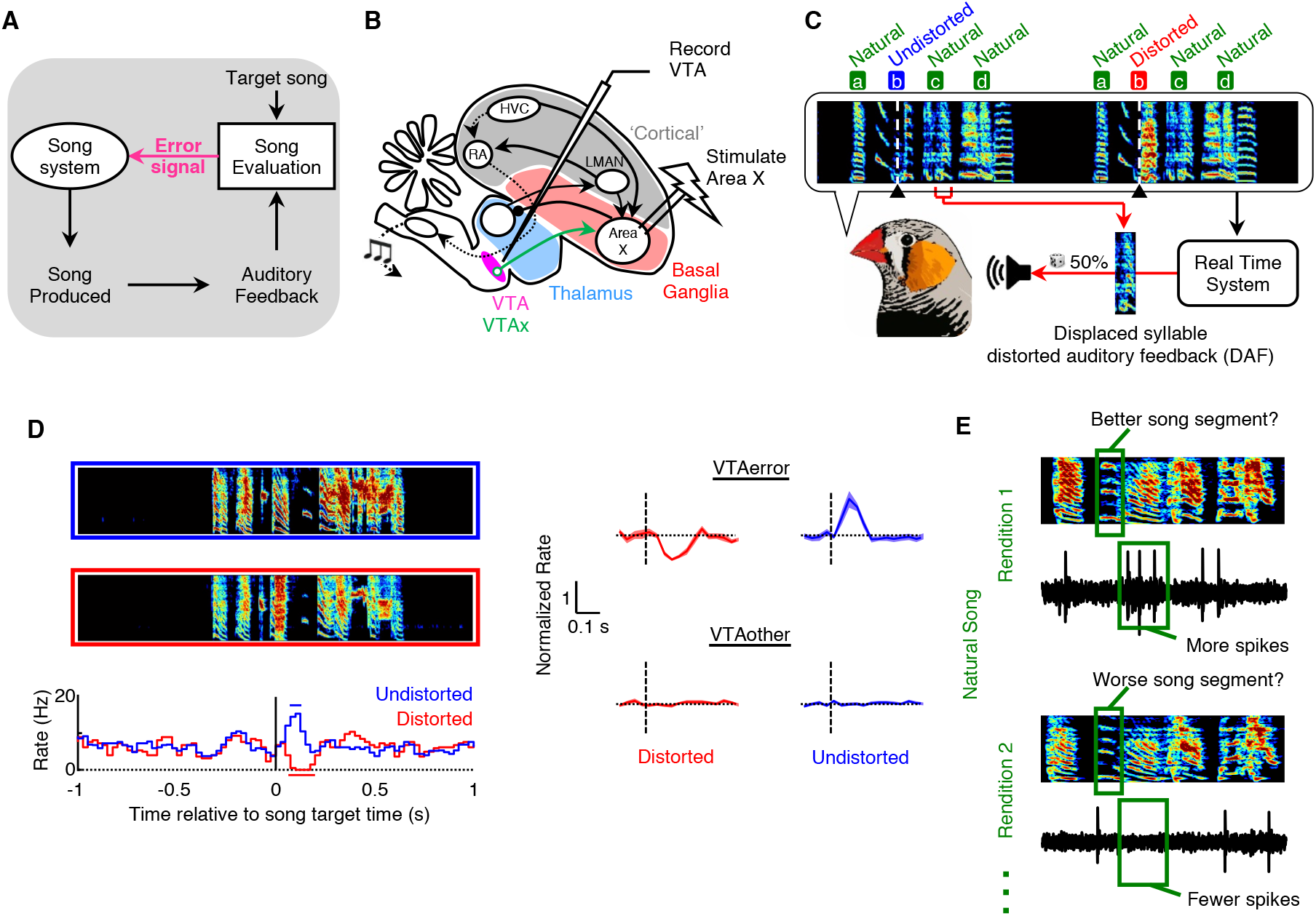
Experimental Identification of Performance Error in VTA DA Neurons in Singing Birds. (A) Evaluation of auditory feedback during singing is thought to produce an error signal for song learning. (B) Basal ganglia (Area X)-projecting DA neurons from VTA were antidromically identified. (C) Example of DAF. The target syllable was randomly distorted across motifs. All other syllables (labeled ‘Natural’) were left undisturbed. (D) Left, top to bottom: example spectrograms of renditions with the target syllable undistorted (enclosed in blue box) and distorted (enclosed in red box); rate histogram of distorted and undistorted renditions (the horizontal bar indicates significant deviations from baseline (p < 0.05, z-test; see STAR Methods)); Right: normalized response to target syllable in VTA_error_ and VTA_other_ neurons (mean +− SEM, see STAR Methods). (E) The experimental results suggest a hypothesis that fluctuations in natural song should also result in VTA_error_ responses.

To test DA’s role in natural behavior, we recorded from DA neurons in the ventral tegmental area (VTA, Figure 1B), and examined how spiking activity correlated with natural song fluctuations (Figure 1E). First, if DA activity following externally distorted and undistorted song (Figure 1D) truly reflects a function of the DA system in performance evaluation, then DA activity should correlate with recent song fluctuations (Figure 1E). Second, DAF-associated error signaling was previously only observed in a small subclass of ‘VTA_error_’ neurons, most of which projected to Area X, the BG nucleus of the song system. ‘VTA_other_’ neurons were defined by the absence of an error signal during singing. We hypothesize that the VTA_error_ population will carry a performance error signal for natural song (Figure 1E), while the VTA_other_ population will not. Thus, we ask in this analysis: do VTA neuron activity patterns relate to fluctuations in natural song? If so, what is the structure of these relationships, and do they relate to a performance evaluation framework, a premotor framework, or both?

To answer these questions, we first parameterized natural song into a low dimensional set of time-varying song features. We then agnostically fit the relationship between rendition-to-rendition variations in song features and spike counts at local time-steps in song across a range of song segment-spike window latencies and identified if and when song feature variations predicted spike counts. Finally, we characterized both the timing and the form of these predictive fits. We find that the activity of the VTA_error_, but not the VTA_other_, neuronal population encodes fluctuations in natural song in a manner consistent with a performance evaluation signal. These results show that basal ganglia-projecting DA neurons may provide continuous evaluation of natural motor performance independent of external rewards or perturbations.

## RESULTS

### A Gaussian Process Model Approach Reveals Song-Spike Relationships

We sought to identify how VTA spiking varied with natural fluctuations in song syllables. To identify relationships between natural song fluctuations and VTA spiking, we chose an eight-dimensional, time-varying representation of song based on established song parameterizations (Figure 2A; see STAR Methods). For each neuron, we identified song syllables and binned both song feature values and spike counts in sliding windows to search for relationships between song fluctuations and spike counts at different latencies (Figure 2B; see STAR methods). We combined all eight song features and binned spike counts into a single multi-dimensional Gaussian Process (GP) regression model (two features shown for illustration) to quantify whether song feature fluctuations predicted spike counts (Figures 2C and S1–S3; see STAR Methods). This strategy flexibly identifies the most relevant dimensions of song variation within a single model. Specifically, we computed an r^2^ value for each model fit using leave-one-out cross validation to assess how well variations in song features predicted spike counts (Figure 2C). Values of r^2^>0 indicates that song feature variations across renditions can predict spike counts; the larger the r^2^ value, the more predictive the song-spike relationship in the model. Finally, we fit the full model to many song-spike latencies and thus built a matrix of r^2^ values for each neuron’s response to song fluctuations, with each r^2^ value in the matrix representing one full model fit (one feature shown for illustration) between a song window-spike window pair (Figure 2D).

**Figure 2.**
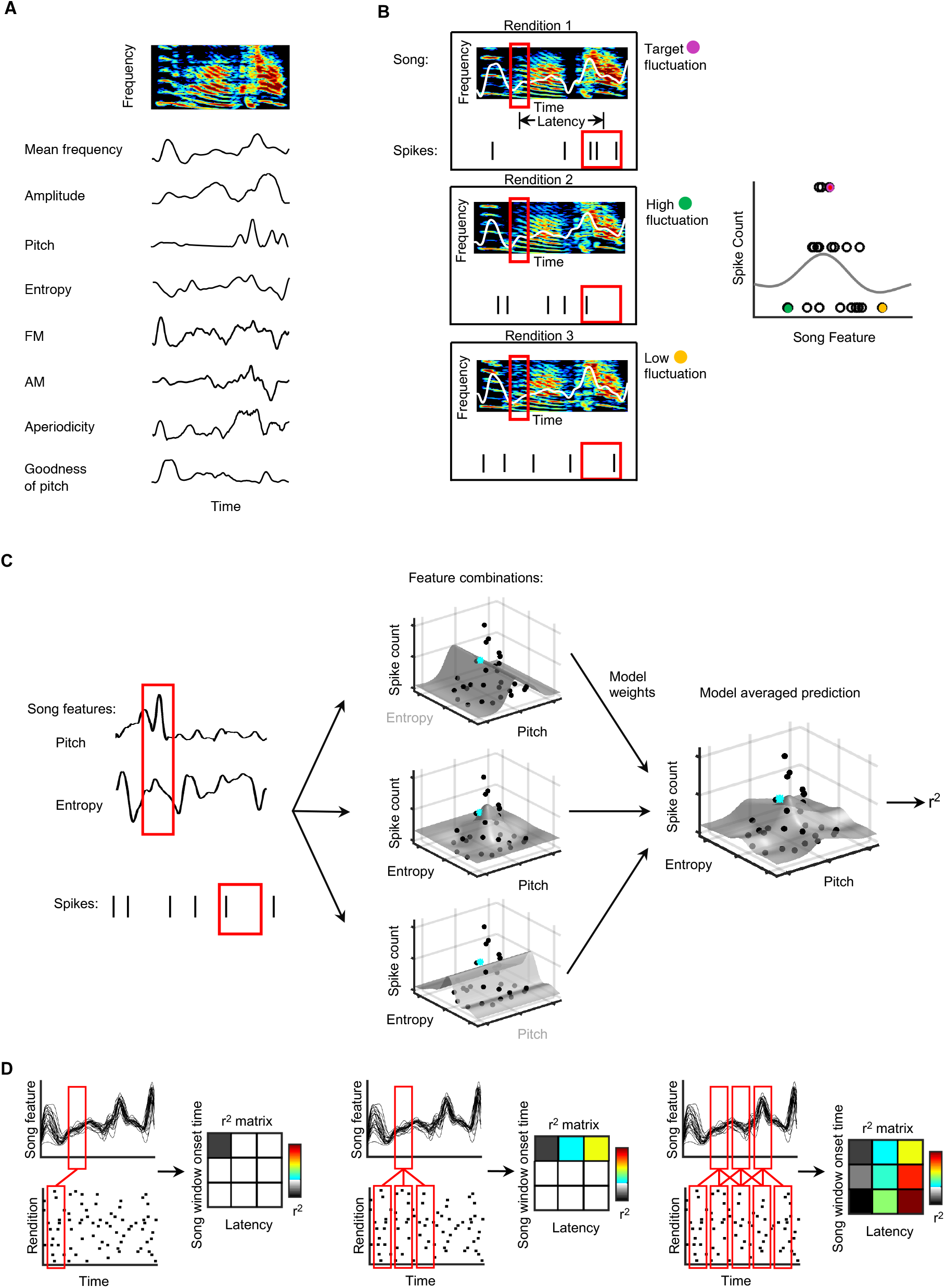
A Gaussian Process Model Approach Reveals Song-Spike Relationships. (A) Natural song was parameterized into eight time-varying song features. (B) Schematic of fitting song fluctuations to spike counts within specific time windows. Local feature averages (one feature shown for illustration) were used to predict local spike counts using a GP model. (C) Schematic of fitting a single, multivariate model using multiple song features. The multidimensional model takes a weighted average of the model predictions from every combination of eight song features (two shown here for illustration). The model’s goodness of fit was quantified by the cross-validated r^2^ (see STAR Methods). (D) The modeling technique shown in (B) and (C) was extended across a range of song windows and song-spike latencies, thus building a matrix of r^2^ values. The top panels show a sliding window along the song (single feature shown for illustration). The bottom panels show the time-aligned spiking activity across renditions in a raster plot. Each entry in the r^2^ matrix (middle panels) represents the fit between one song window and one spike window, shown here connected with red lines.

### Timing of Song-Spike Relationships for VTAerror Neurons is Consistent with an Evaluative Process

Using the GP model approach described above, we asked if significant relationships between natural song fluctuations and VTA neuron spiking exist, and if so, at what song-spike latencies they occur. If VTA spiking is predictive of upcoming syllable fluctuations in a premotor fashion, then significant relationships would be observed at negative lags. Alternatively, if VTA spiking is playing an evaluative function, then variations in spike counts should follow variations in syllable acoustic structure, and relationships should be observed at positive lags. Based on past work (Gadagkar et al., 2016), an evaluation signal is predicted to occur at a positive lag of ~50 ms with a duration range of 0 to 150 ms. Figure 3A shows an example VTA_error_ neuron’s song-spike relationship (the r^2^ matrix) for a single syllable. The y-axis is the midpoint of each song window (song window width = 35 ms) aligned to syllable onset (t=0). The x-axis is the latency, defined as the time between the song window midpoint and the spike window midpoint (spike window width = 100 ms). Colored pixels in the r^2^ matrix indicate that song feature fluctuations predict spike counts (r^2^ > 0); greyscale pixels indicate that song feature fluctuations do not predict spike counts (r^2^ <= 0). The pink box indicates the song-spike latencies (0-150 ms) where we expect to see evaluation-like relationships based on the DAF experimental results (Figure 1D). We assessed the significance of finding predictive fits by shuffling entire spike trains relative to song renditions and refitting our model across all latencies and song windows (Figures 3A bottom and S4; see STAR Methods). This population method of shuffling the data preserves the underlying temporal correlation structure of song and spiking while randomizing the song-spike relationship and allowed us to assess the significance of the entire set of model fits and account for multiple comparisons (see STAR Methods). The bottom matrix in Figure 3A shows the r^2^ values from one such randomized shuffle of the same neuron’s activity patterns. Positive r^2^ values were found to be less frequent in the shuffled data (one-sided z-test: p < 0.01).

**Figure 3.**
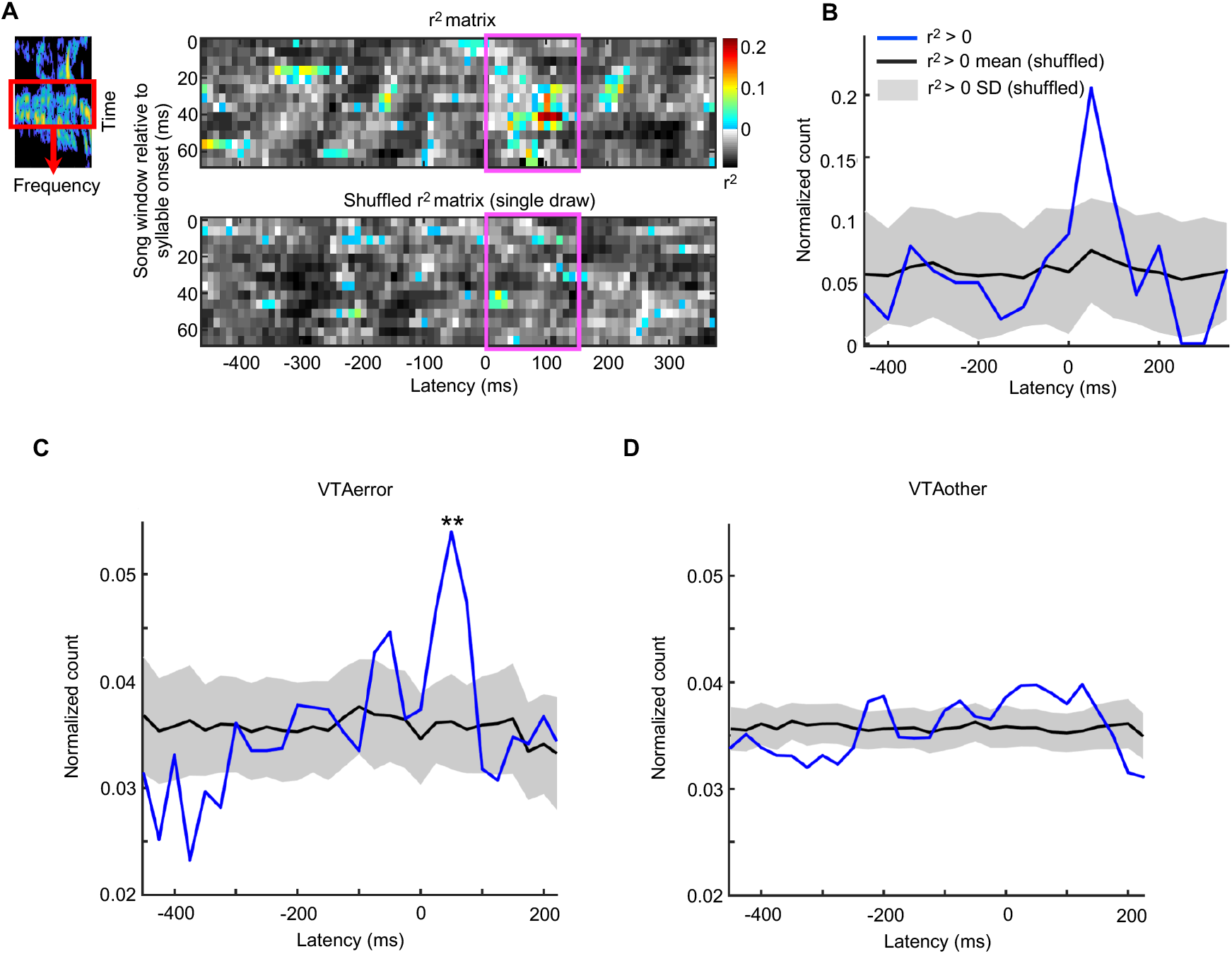
Timing of Song-Spike Relationships for VTA_error_ Neurons Suggests an Evaluative Process. (A) Spectrogram of example syllable (top left). Heat map of r^2^ values for fitted relationships between local song feature averages and binned spike counts (top right). r^2^ > 0 indicates a predictive relationship. The pink box indicates the region where the latency matches the hypothesized response for a PPE, 0-150 ms. The lower heat map shows an r^2^ matrix for a shuffled version of the data (see STAR Methods). (B) Histogram of latencies for predictive fits shown in (A). (C) Latency distribution of predictive fits over all VTA_error_ neurons (N=19) showed a significant peak in the number of responses in the expected PPE time window (** indicates p < 0.01, see STAR Methods). (D) Same as in (C), but for the VTA_other_ neuron population (N=23).

We then analyzed the temporal relationship between song features and spiking by comparing the latency distribution for all song window-spike window pairs within the r^2^ matrix in which song features predicted spike counts (r^2^ > 0) to the latency distribution for pairs without predictive relationships (r^2^ <= 0) (Figure 3B). The latencies of the predictive fits were clustered within the expected error evaluation range (0-150 ms) (Figure 3B). Figure 3C shows the result of the same analysis performed across all the VTA_error_ neurons in our dataset (N=19). In Figure 3C, the blue line indicates the song-spike latency distribution for which song fluctuations predict spike counts (r^2^ > 0) while the black line and grey shading are the mean and standard deviation of the same latency distributions across all randomized population shuffles. The true data showed a large peak within the expected PPE latency range (3.74 standard deviations from the mean, one-sided z-test: p < 0.01; see STAR Methods). We found that song fluctuations are most predictive of spike counts 0-100 ms after the song fluctuation occurs, consistent with a PPE-like signal based on our previous DAF experiments (Figure 1D). In addition, across the population of VTA_error_ neurons there were significantly more predictive fits within the PPE latency window than expected by chance (p <= 0.01; see STAR Methods). Thus, the timing and frequency of the predictive song-spike relationships was remarkably consistent with a PPE-like response to natural song variations.

We next performed the identical analysis on a population of VTA_other_ neurons (N=23), which did not show an error-like response in previous DAF experiments (Figure 1D). The number of predictive song-spike relationships from this population was also significantly larger than expected by chance (bootstrap test, p <= 0.01, see STAR Methods). However, unlike the VTA_error_ neurons, the predictive relationships from this population did not cluster within the PPE latency range, nor did the variance of the distribution significantly deviate from the randomized latency distributions (p=0.21; Figure 3D; see STAR Methods). Thus, consistent with results from the DAF experiments (Figure 1D), only the responses of VTA_error_, and not the non-error responsive VTA_other_, neuron population were predicted by natural song fluctuations within the expected PPE latency range with significantly increased frequency. The same neurons that exhibited error responses to the DAF sound exhibited significant relationships with natural syllable fluctuations. Remarkably, both the DAF-induced error and the natural fluctuations were at a similar latency with respect to song.

### The Form of the Predominant Song-Spike Relationship for VTA_error_ Neurons is Consistent with Song Maintenance

The hypothesis that VTA_error_ neurons evaluate natural song fluctuations led to further predictions about the forms of these song-spike relationships. If a bird is trying to maintain the acoustic structure of a syllable, then typical variations should be followed by more spikes and rare, outlying syllable variations should be followed by fewer spikes (Figure 4A, top left). Alternatively, if the bird is trying to modify a syllable, e.g. increase its pitch, then the relationship between syllable acoustic structure and spike should be directional: spike counts should peak at whatever shifted variant to which the bird aspires but is not yet consistently producing (Figure 4A, right panels). We did not expect PPE-like signals to have multiple maxima in a *disruptive* fit: we assumed there is a single ‘best’ version of the song at each time-step (Figure 4A, bottom left).

**Figure 4.**
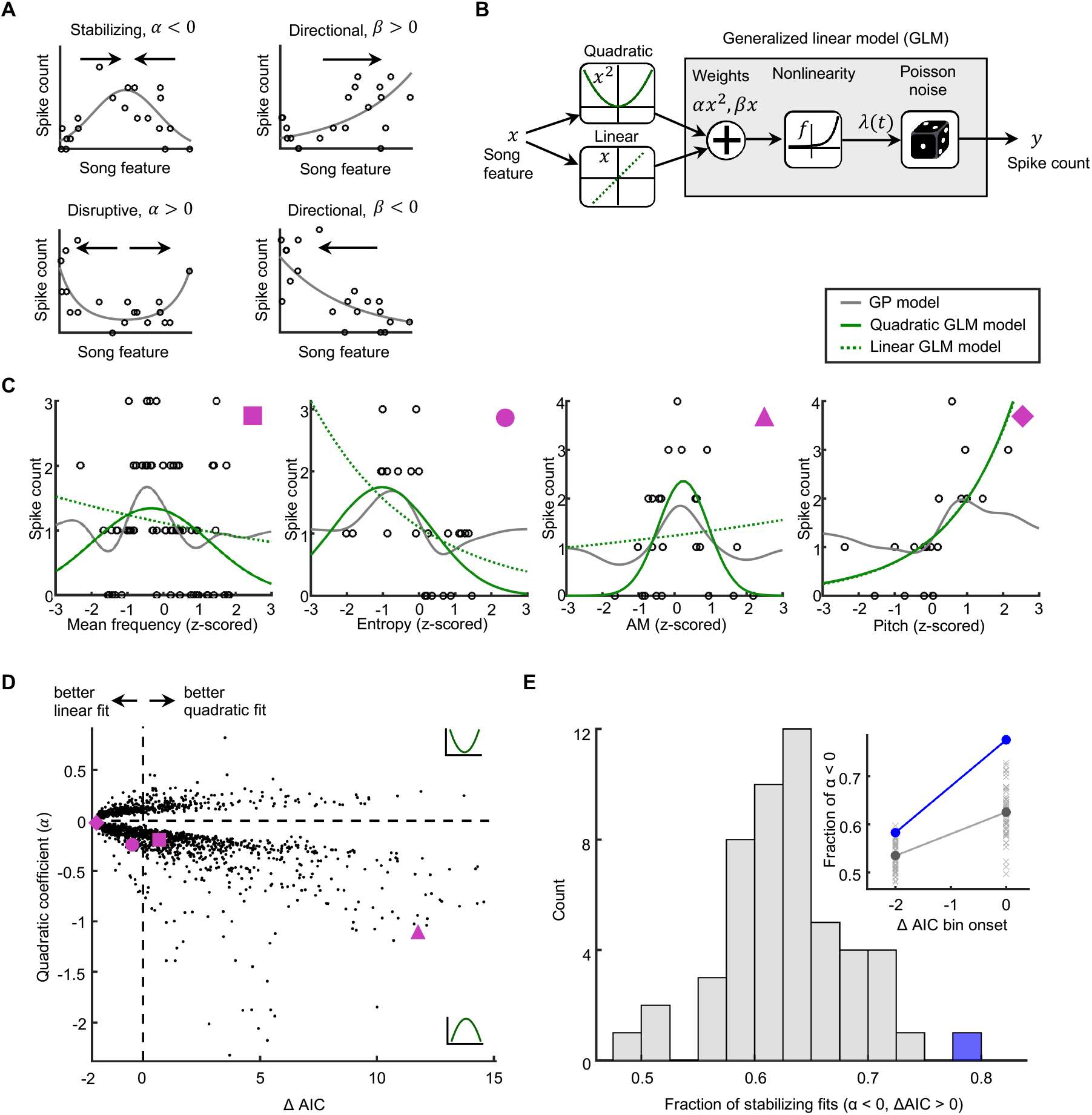
The Form of the Predominant Song-Spike Relationship for VTA_error_ neurons is Consistent with Song Maintenance. (A) The form of a song-spike relationship determines how the song is being reinforced. α and β correspond to the quadratic and linear coefficients in the GLM shown in (B). (B) Schematic of the nested GLM fitting process to quantify tuning curve shape for VTA_error_ neuron activity to natural song fluctuations. (C) Example tuning curves obtained with the GP model, l-GLM, and q-GLM between single song features and spike counts for a selection of song-spike model fits. Each point on each plot represents a single rendition. Pink shapes denote fit locations marked in (D). (D) The quadratic coefficient for all q-GLM model fits to predictive song-spike relationships (defined within the GP model) as a function of ΔAIC values in the GLM model comparison within the PPE latency range. Pink shapes denote fits shown in (C). (E) Fraction of stabilizing fits (negative quadratic coefficient) for all fits better described as quadratic than linear (ΔAIC > 0) compared to shuffled population fractions. The blue point is the data and each value in the gray histogram is a single fraction from an independent population shuffle (see STAR Methods). The data showed a greater fraction of stabilizing fits than expected by chance (p-value < 0.01). Inset: same distribution but now shown for both the binned ΔAIC > 0 group and Δ*AIC* ∈ [−2, 0]. The blue point is the true fraction and gray points are fractions from shuffled populations (see STAR Methods).

To test these possible outcomes, we characterized tuning curve shapes of song-spike relationships. For this analysis, we focused our attention on the subset of GP model fits that were predictive (r^2^ > 0). Within this subset, we further selected single-feature fits that were also predictive (r^2^ > 0 for the 1D feature fit). We selected this subset of song-spike relationships because we are interested only in the tuning curves that might actually carry information about song. We re-fit all such song-spike relationships with a generalized linear model (GLM) using both linear (l-GLM) and quadratic (q-GLM) features (Figure 4B; see STAR Methods). We chose these models because the parameters can be used to directly quantify aspects of the tuning curve shapes. If the song-spike fit has a single peak, the spiking response is stabilizing, and the quadratic coefficient of the q-GLM is negative (Figure 4A, top left). If the song-spike fit has two peaks, the spiking response is disruptive, and the quadratic coefficient of the q-GLM is positive (Figure 4A, bottom left). If the song-spike fit is monotonic, the spiking response is directional, and the l-GLM (with only linear features) is the more appropriate model (Figure 4A, right panels). Figure 4C shows examples of predictive relationships between individual song features and spike counts along with all model fits (GP, q-GLM, and l-GLM). Each point on these plots represents the song feature value and the spike count for a single rendition. Specific models were better fits for some distributions than others. For example, in panel 4 (Figure 4C, fourth panel from left), the q-GLM produced the same model fit as the l-GLM because the quadratic term added no improvement to the fit, whereas in panel 3 (Figure 4C, third panel from left), the quadratic term was necessary to accurately follow the spiking response and thus the q-GLM resulted in a better fit than the l-GLM.

When the spiking response is either clearly stabilizing or disruptive, the quadratic coefficient of the q-GLM distinguishes between these two response types making the q-GLM the better model choice. We used the Akaike Information Criterion (ΔAIC) to compare the relative success of the q-GLM and the l-GLM (see STAR Methods). ΔAIC > 0 indicates that the quadratic model outperforms the linear model, considering both the likelihood of the model fit and the complexity of the model used. ΔAIC = −2 indicates the quadratic model provides no benefit over the linear model. The larger the ΔAIC, the better the q-GLM fit relative to the l-GLM. In Figure 4D, each point represents a fit to a single song feature with an r^2^>0 within a multi-dimensional model fit with an r^2^>0 for the population of VTA_error_ neurons within the expected PPE latency range. We found a greater fraction (0.78) of q-GLM fits with negative quadratic coefficients, which indicates more stabilizing tuning curves than disruptive tuning curves (Figure 4D). Furthermore, when the quadratic model outperformed the linear model, the fraction of stabilizing fits also increased (Figure 4E). Thus, the predictive fits from the GP model had significantly more stabilizing tuning curves than disruptive when their shape was better characterized as quadratic rather than linear, as we expect for a PPE signal with a single best outcome (2-tailed z-test: p < 0.02). This finding is consistent with our hypothesis that a PPE signal should respond most strongly to a single best performance of song. The ΔAIC measure also allowed us to examine the fraction of tuning curves that are better fit by a linear versus quadratic model. The VTA_error_ population did not differ from chance in this fraction (fraction fits with ΔAIC > 0 = 0.43; 2-tailed z-test, p = 0.34), consistent with a PPE signal with both directional and stabilizing responses depending on the current level of song error.

## DISCUSSION

Value judgements in the brain are necessary to drive appropriate changes in behavior during learning. Using experimentally constrained tasks with external rewards, previous studies have found that DA neurons in VTA can encode a key component of value judgement: the mismatch between expected and actual reward outcomes, the reward prediction error (Schultz *et al.*, 1997). However, extending these findings to natural behavior and intrinsic reward has been a challenge. Here, we made use of a novel opportunity to use an experimental context to partition songbird VTA neurons into error and non-error classes and analyze their spiking responses in the context of a natural behavior (Gadagkar et al., 2016). We compared natural song fluctuations at a local, within-syllable scale to variations in spike counts of VTA neurons. We developed a Gaussian Process regression analysis to quantify the non-stationary spiking response to variations in performance at different points in song and with different temporal relationships to song. We found evidence that VTA DA neurons’ activity patterns correlate with variations in natural song in a manner consistent with performance evaluation: both the timing and tuning properties of the DA response was consistent with a PPE-like response. This finding corroborates and extends complementary discoveries of RPE signals emerging from mammalian DA neurons in VTA in experimentally imposed tasks. We did not find significant temporal relationships between DA and song fluctuations consistent with a premotor signal as has been observed in previous studies of DA (Barter et al., 2015; Engelhard et al., 2019). Our results are the first direct evidence we are aware of that DA neurons in VTA respond to fluctuations in natural behavior in a manner consistent with evaluation.

Two important predictions from these PPE-like signals we find in DA VTA neurons is that, one, future song renditions will shift towards song variations which correlate with the peak response in the DA neurons and that, two, this shift will be accompanied by a decrease in the PPE peak response. We could not address these predictions here because we analyzed single recording sessions of limited duration. This will be an important direction for future work and will help disambiguate the role of the VTA DA responses from other possible relationships to song. We chose a pre-defined set of song features (N=8), which have been shown to represent biologically relevant song variations in previous studies, because we focused on single sessions with limited data. Future work could apply more flexible, non-parametric dimensionality reduction methods using more song renditions to better identify VTA’s relationship to song features that are most modulated by the bird at different points in song (Goffinet et al., 2021; Kollmorgen et al., 2020).

While, as a population, the non-error VTA_other_ neuron activity did not relate to song fluctuations in a manner consistent with a PPE signal, many neurons exhibited correlated relationships to song (Chen et al., 2021). These findings are consistent with previous studies in mammals which found that both DA and non-DA neurons in VTA contribute to an RPE calculation and that elements of the RPE signal are computed, in part, locally within VTA (Cohen et al., 2012; Dobi et al., 2010; Ju Tian, 2016; Wood et al., 2017). Correlations with song variations in this population could represent components of the PPE calculation.

This project uses the structure of an experimentally grounded characterization of individual neurons’ response to song-triggered, distorted auditory feedback to analyze the same neurons during natural behavior. The connection to an existing experiment (Gadagkar et al., 2016) as well as to a Reinforcement Learning framework (Sutton and Barto, 1998) anchors our interpretations of natural behavior in a constrained laboratory paradigm and theory. The unusually high stereotypy of the natural behavior we consider, zebra finch song, allows reasonable inferences to be made both in the experimental and natural context about the behavior of the bird and a reasonable way to characterize and align a complex, natural behavior. We found a parallel relationship, including a striking temporal correspondence, between the VTA_error_ neuron activity in experimental and natural contexts that corroborates the experimental finding that VTA_error_ neurons encode time-step specific performance prediction errors in song. Our analysis of natural song addresses the critique that the DAF experimental paradigm is aversive rather than perturbative and thus qualitatively different from natural song evaluation. A frequent debate in neuroscience is whether artificial behavioral paradigms serve as true building blocks for understanding neural activity in complex, freely behaving contexts, or whether they represent a different, overly-simplified context that will not extrapolate to natural behavior. This experimentally guided study of natural behavior is a fruitful direction that permits the control of experimental contexts and the complexity of natural contexts to interact and build upon one another.

## ACKNOWLEDGMENTS

This work was supported by NIH/NINDS (grant# R01NS094667), Pew Charitable Trusts, and the Klingenstein Neuroscience Foundation (J.H.G.), a Simons Collaboration on the Global Brain (SCGB) grant (A.L.F and A.D.), an SCGB Postdoctoral Fellowship and NIH/NINDS Pathway to Independence (grant# K99NS102520 and R00NS102520) award (V.G.).

## AUTHOR CONTRIBUTIONS

Conceptualization, A.D., K.W.L., J.H.G., A.L.F., and V.G.; Experimental Methodology and Investigation: J.H.G and V.G. Analysis Methodology, A.D., K.W.L., A.L.F., and V.G.; Software and Formal Analysis: A.D., K.W.L., and A.L.F.; Writing-Original Draft, A.D. and V.G.; Writing-Review and Editing, A.D., K.W.L, J.H.G., A.L.F., and V.G.; Funding Acquisition, J.H.G., A.L.F, and V.G.; Supervision, J.H.G., A.L.F, and V.G.

## STAR METHODS

### RESOURCE AVAILABILITY

#### Lead Contact

Further information and requests for resources should be directed to and will be fulfilled by Vikram Gadagkar.

#### Materials Availability

This study did not generate new unique reagents.

#### Data and Code Availability

The data and code generated during this study are available at (to be created code repository).

### EXPERIMENTAL MODEL AND SUBJECT DETAILS

Subjects were 26 adult male zebra finches 74-300 days old singing undirected song. All experiments were carried out in accordance with NIH guidelines and were approved by the Cornell Institutional Animal Care and Use Committee. During implant surgeries, each bird was anesthetized with isoflurane and a bipolar stimulation electrode was implanted into Area X at established coordinates (+5.6A, +1.5L relative to lambda and 2.65 ventral relative to pial surface; head angle 20 degrees). Intraoperatively in each bird, antidromic methods were used to identify the precise part of VTA containing VTAx neurons. Next, custom made, plastic printed microdrives carrying an accelerometer, linear actuator, and homemade electrode arrays (5 electrodes, 3-5 MOhms, microprobes.com) were implanted into this region.

### METHOD DETAILS

#### Syllable-targeted distorted auditory feedback

Detailed description of all aspects of the distorted auditory feedback (DAF) experiments is described elsewhere (Gadagkar et al., 2016). Descriptions of experimental details relevant to this study are presented here. Postoperative birds were placed in a sound isolation chamber equipped with a microphone and two speakers which provided DAF. To implement targeted DAF, the microphone signal was analyzed every 2.5 ms using custom Labview software. Specific syllables were targeted either by detecting a unique spectral feature in the previous syllable (using Butterworth band-pass filters) or by detecting a unique inter-onset interval (onset time of previous syllable to onset time of target syllable) using the sound amplitude as previously described (Ali et al., 2013; Hamaguchi et al., 2014; Tumer and Brainard, 2007). In both cases a delay ranging from 10-200 ms was applied between the detected song segment and the target time.

To ensure that DAF would not be perceived as an aversive stimulus, the DAF sound had the same amplitude and spectral content as normal zebra finch song. For broadband DAF (n = 16 birds), DAF was a broadband sound band passed at 1.5-8kHz, the same spectral range of zebra finch song (Andalman and Fee, 2009). For displaced syllable DAF (n = 10 birds), DAF was a segment of one of the bird’s own syllables displaced in time. For both types of DAF, the amplitude was carefully measured with a decibel meter (CEM DT-85A) and maintained at less than 90 dB, the average peak loudness of zebra finch song (Mandelblat-Cerf et al., 2014). This ensured that DAF was not a particularly loud sound for the bird; the distorted part of the song was significantly softer than the loudest parts of the song.

Target time in the song was defined as the median time of DAF onset across target syllables; jitter of the target time was defined in each bird as the standard deviation of the distribution of DAF onset times relative to the target syllable onset. Syllable truncations following DAF were rare and were excluded from analysis.

#### Electrophysiology

Neural signals were band pass filtered (0.25-15 kHz) in homemade analog circuits and acquired at 40 kHz using custom Matlab software. Single units were identified as Area X-projecting (VTAx) by antidromic identification (stimulation intensities 50-400 μA, 200 μs on the bipolar stimulation electrode in Area X). All neurons identified as VTAx were further validated by antidromic collision testing. Spike widths were computed as the trough-to-peak interval in the mean spike waveform.

#### Spike sorting and analyzing responses to distorted auditory feedback

Spike sorting was performed offline using custom Matlab software. Firing rate histograms were constructed with 25 ms bins and smoothed with a 3-bin moving average. To calculate the significance of error responses (Figure 1D), spiking activity within ±1 second relative to target onset was binned in a moving window of 30 ms with a step size of 2 ms. Each bin after the target time was tested against all the bins in the previous 1 second (the prior) using a z-test (Mandelblat-Cerf *et al.*, 2014). Response onset (latency) was defined as the first bin for which the next 3 consecutive bins (6 ms) were significantly different from the prior activity (z-test, p < 0.05); response offset was defined as the first bin after response onset for which the next 7 consecutive bins (14 ms) did not differ from the prior (p > 0.05, z-test); the response onset and offset were required to bracket the maximum (undistorted) or minimum (distorted) response after target time.

#### Parameterizing song

We used open source MATLAB software, Sound Analysis Pro 2011 (SAP 2011), to assemble the spectrogram as well as to define and extract song features. SAP 2011 is a customized software package written to analyze animal communication and is originally and most frequently used to study bird song (Tchernichovski et al., 2000). We used an existing SAP feature set for our parameterization because these features have been used in many previous studies to link zebra finch song variations to neural activity or neuromodulator concentrations (Kao et al., 2005; Leblois et al., 2010; Woolley and Kao, 2014), to study variation in song over development (Deregnaucourt et al., 2004; Lipkind and Tchernichovski, 2011; Ravbar et al., 2012) and to drive adult learning in DAF paradigms (Andalman and Fee, 2009; Sober and Brainard, 2009; Tumer and Brainard, 2007). Therefore, we can use this form of dimensionality reduction of song knowing in advance that these dimensions are behaviorally relevant to song variation in other contexts. The features extracted were Wiener entropy, pitch, goodness of pitch, amplitude, amplitude modulation (AM), frequency modulation (FM), mean frequency, and aperiodicity. These features result in an eight-dimensional representation of song at each time-step. We further applied a moving-average filter (35 ms) to smooth the feature signals in time and sampled the smoothed value every 5 ms across song.

#### Aligning syllables across renditions

To compare song across renditions, syllables were classified using custom Matlab code (Gadagkar et al., 2016). Clusters of unique syllables were labelled alphabetically as ‘a’, ‘b’, ‘c’ etc. depending on order within a rendition. The number of syllables each bird sings varies bird-to-bird from 3-7 syllables. We identified syllable onsets and offsets across renditions for every syllable set in which there were greater than 15 renditions of that syllable using an amplitude threshold chosen to match the amplitude variance of that syllable. All alignments were further checked by eye. Renditions in which alignment was ambiguous by eye were excluded from analysis.

All syllable types (i.e. ‘a’ or ‘b’ etc.) were isolated and aligned across renditions by syllable onset times. Individual syllable types have a stereotyped, characteristic duration; however, there is some variation of this duration from rendition-to-rendition. In order to make sure that minor differences in syllable lengths were not misaligning local syllable features at the later parts of the syllable, we linearly time-warped the feature wave forms of each syllable rendition such that they all lasted the median duration of that syllable type (Kao et al., 2008).

#### Parameterizing and aligning spiking activity

Spike sorting was performed offline using custom MATLAB software (Gadagkar et al., 2016). For every syllable, we considered the spike train ± 500 ms around the syllable onset. In order to align spiking activity to the song features, we first applied the same linear time-warping map to the spike train that we used to align syllables for each rendition (Kao *et al.*, 2008). In all cases, we applied this map to the time window in which the syllable took place. When possible, we generated a piece-wise linear time warping map based on syllable boundaries in surrounding syllables. In the time windows where there was no song with which to build a time warping map, we left the spike train un-warped.

We binned spike counts within a sliding window (100 ms) across the 1000 ms length of spike train we considered for each syllable. We chose this spike count window based on the firing rate of the VTA-error neurons we considered (mean firing rate = 13 ± 5 Hz).

#### Fitting spikes to song with a Gaussian process regression

The goal of our analysis is to quantify non-stationary spiking responses to a time-varying sensory signal, with the following characteristics. First, if VTA neuron activity encodes prediction error responses to song fluctuations, these responses would be specific to the time in song; an identical vocalization occurring at the beginning of the song might elicit a very different response than at the middle. Second, the relevant dimensions of the signal space could vary throughout the song; thus, different parameterizations of the song might provide better low-dimensional representations of error-relevant song variation at different song time-steps. Third, the form of a PPE-like tuning curve could also vary across the song.

Towards this goal, we used a regression approach to determine if spike counts are related to the variation in song. The relationship between spike counts and song is likely non-linear and related to a variable number of features depending on the point in song. To address this, we used a non-parametric Gaussian process (GP) regression to fit the relationship between our eight song features and spike counts within single time windows (Williams, 2006).

There are multiple sources of model uncertainty in this task: it is unclear which and how many features to use at a given point in song. Furthermore, the prediction of the model depends heavily on which features are used. To address this uncertainty, we used a Bayesian model averaging approach to determine the predicted spike count wherein we integrated over all possible feature combinations and weighted their predictions according to their posterior probability given the observed spike counts (Hoeting, 1998).

For each neuron, in every non-target (distorted or undistorted) syllable for which there were *N* ≥ 15 renditions, we sampled the smoothed song features every 5 ms across the syllable and sampled spike counts in 100 ms windows every 10 ms across 1s of spike train centered around syllable onset. We fit the multi-dimensional GP model across all song segment-spike bin pairs and generated song-spike relationships at many time latencies. We additionally fit a GP model using each feature individually. For all of these fits we computed the *r*^2^ value.

#### Construction of the Gaussian process regression model

We modeled the relationship between the set of N z-scored song features on a single rendition i, ***x***_*i*_, and the spike counts on that given rendition, *y*_*i*_, in single time windows (e.g. the song feature values 20 ms after syllable onset and the spike count in a 100 ms window, 75 ms after syllable onset). To address this, we used a non-parametric Gaussian process (GP) regression to fit the relationship between the eight song features and spike counts across song and spike window pairs (Williams, 2006). We used a Bayesian model averaging approach to combine a weighted average of GP regressions using all subsets of song features into a single model prediction.

We selected a subset of features, M, for a single GP regression, where feature is indexed by *i* = 1, 2, …, *N* such that 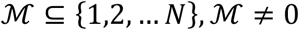, N = 8. The GP regression model for a single set of 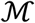 is:

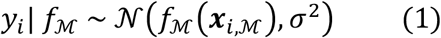

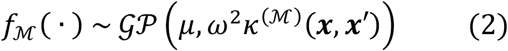

where f is a function relating song features to spike rate and *κ* is the covariance function and defines how spike counts will be correlated with one another in feature space. We used the commonly selected kernel function for *κ*,

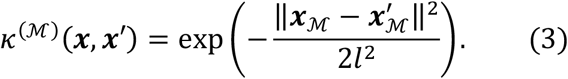

*ω*^2^ is the GP variance term and specifies how strongly the spike counts vary as a function of the song features. σ^2^ is the variance term that captures noise in the spike count (i.e. how much the spike counts vary at a single point in song feature space). The length scale, *l*, determines how close points must be in feature space to have correlated spike counts. To reduce computational complexity, we set *l* = 0.5, for all model fits and z-scored the individual song features at each time step we considered.

Although we do not observe f directly, the GP framework allows us to compute the marginal likelihood of the data with respect to the model parameters. The likelihood of all *T* renditions of spike count-song segment pairs is:

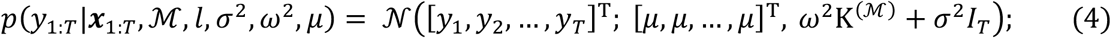

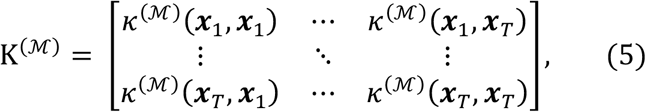

where *I*_*T*_ is the identity matrix of dimension *T*.

The prediction mean-squared error for the GP model is:

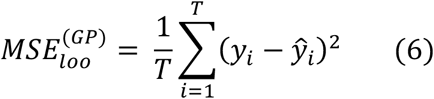

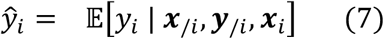

where 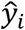 is the predicted spike count from the model for rendition *i*, and ***x***_/*i*_ and ***y***_/*i*_ are the song features and spike counts for all renditions except for the *i*^*th*^ rendition.

We determined the predicted spike count by applying a Bayesian model averaging approach. We integrated over all possible values of 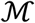 and then weighted their predictions based on their posterior probability given observed spike counts (Hoeting, 1998):

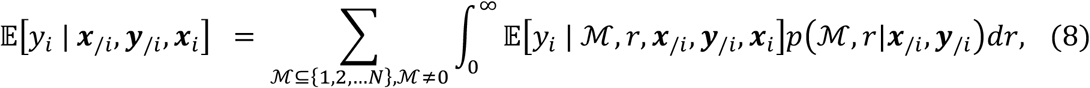

where 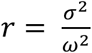 is the ratio of the GP variance to the observation noise. We integrated over all possible values of 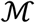 and weighted their predictions according to their posterior probability given the observed spike counts.

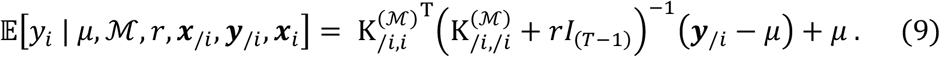

Thus, we incorporated all combinations of features into a single model prediction for each song-spike count pair. We re-parameterized (σ^2^, *ω*^2^) to (*ψ*^2^, *r*^2^) where *ψ*^2^ is the total variance:

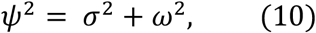

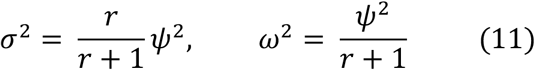

And evaluated the posterior over model parameters using Bayes’ rule:

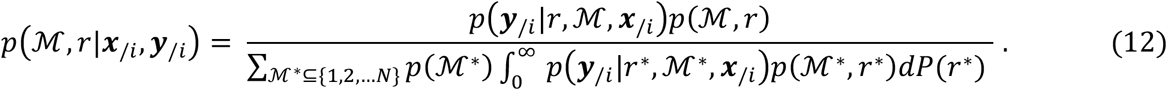

We again used Bayes’ rule to compute the likelihood term in Eq. 12:

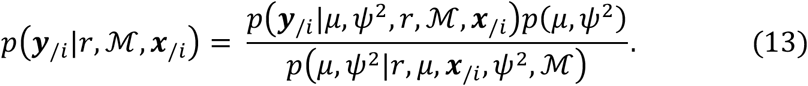

The likelihood term is computed as in Eq. 4. We again used Bayes’ rule to compute the posterior over *μ* and *ψ*^2^:

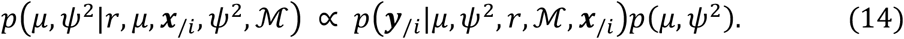

We then placed a conjugate normal-inverse gamma prior over *μ* and *ψ*^2^:

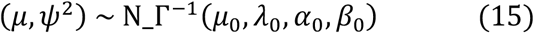

where,

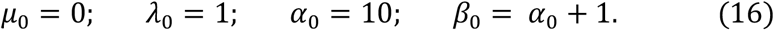

Thus,

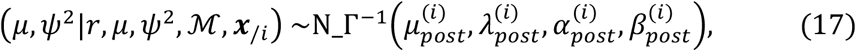

where,

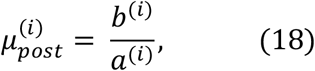

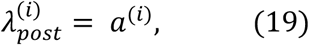

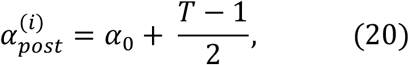

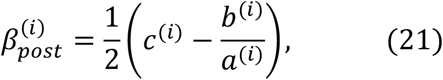

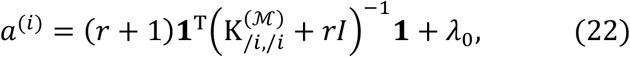

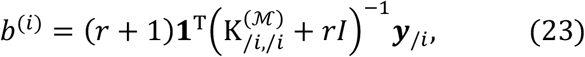

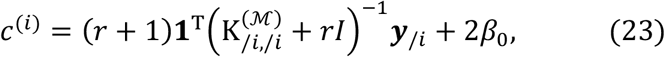

where **1** is a vector of ones. With this, we can compute all of the terms in Eq. 13.

The integral over Eq. 12 is over one dimension and thus tractable to compute. We chose a discrete distribution for the prior *P*(*r*) to increase computation speed:

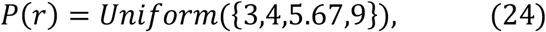

such that the GP model could account for 25%, 20%, 15% or 10% of the total variance.

We imposed a truncated binomial prior over the number of included features such that 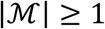, that favored models with fewer features (i.e., sparse models):

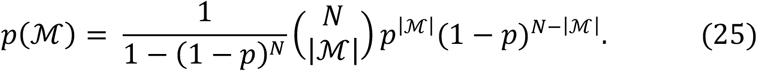

We set p=0.1 so that approximately 2/3 of the prior probability mass rests on single-feature models. We could integrate over a sparse prior in our model, rather than a shrinkage prior such as the Lasso, because we considered only a small (N=8) set of features (Park and Casella, 2012). Using this normal inverse-gamma description of the posterior, we can compute the prediction of *y*_*i*_, given 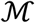 and *r*:

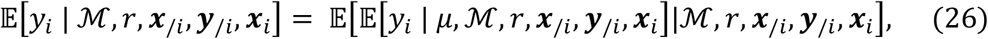

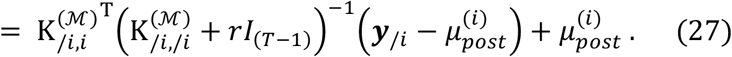

We then insert Eq. 27 and Eq. 12 into Eq. 8 to obtain the prediction of *y*_*i*_.

#### Construction of the latency distribution

We defined the latency distribution as the set of all latencies between spike bins and song feature windows in which there was a predictive relationship (r^2^>0) within the GP model.

#### Characterizing tuning curves of cell responses

The GP model is flexible in that it will fit any relationship between the independent and dependent variables and is computationally efficient. However, from the output of the model we have no easily interpretable means of characterizing the shape of the fit. In order to characterize the form of the spike-count to song relationships across the large number of fits we assessed, we needed an automated way to categorize the shapes of the tuning curves.

To do this, we used a generalized linear model (GLM) with (q-GLM) and without (l-GLM) a quadratic transformation of the song features (Il Memmming Park, 2013). A GLM consists of a stimulus filter, an invertible non-linearity (the link function) and a stochastic exponential non-linearity, such as a Poisson process:

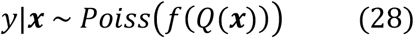

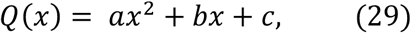

where y is the spike count, f is the inverse link function, x is the stimulus and Q is a quadratic function of *x* with coefficients *a*, *b*, *c*. We considered song features individually in this tuning curve analysis, so dim(***x***) = 1, and the quadratic coefficients became scalars (a, b, c). We took the link function to be an exponential and the noise process to be Poisson. Thus, we can fit the quadratic coefficients by maximizing the log-likelihood:

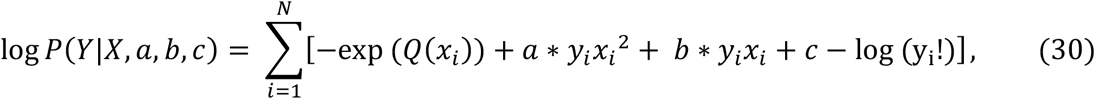

We maximized the log-likelihood numerically using conjugate gradient methods. The sign of the quadratic coefficient, a, of this model determines whether the data is better fit by an upwards-facing, quadratic basis in which the data is double-peaked, or a downwards-facing quadratic basis in which the data is single-peaked. We compared this model to a nested model fit where the quadratic term is set to zero (l-GLM).

We compared the performance of the two models using the Akaike information criterion (AIC) (Akaike, 1974). The AIC metric is defined as:

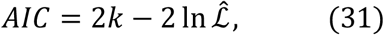

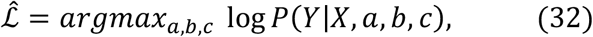

where 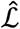 is the maximum of the log-likelihood function for a given model and *k* is the number of estimated parameters. This metric balances goodness of fit with model complexity. A lower AIC metric indicates better performance. Therefore, the difference in the AIC metrics of two models indicates the relative success of one model over another, taking into account differences in model complexity (Akaike, 1974; Raftery, 1995). We can then ask, when the quadratic model (q-GLM) is a better fit to the data than the linear model (l-GLM), does the tuning curve relationship of spike counts to song features show a positive or negative curvature? We predicted that a PPE-like signal should have a negative curvature.

We compared the q-GLM and l-GLM models on all GP model fits with *r*^2^ > 0 for all individual song features which had themselves predictive fits within the multi-dimensional model. We calculated the fraction of fits with the quadratic coefficient, a < 0, as a function of the AIC comparison between the two models:

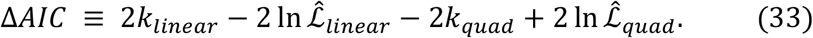

### QUANTIFICATION AND STATISTICAL ANALYSIS

#### Evaluating the Gaussian Process model performance

To evaluate GP model performance we use a leave-one-out cross validation method to estimate the mean-squared prediction error for new observations as in (Vehtari, 2017):

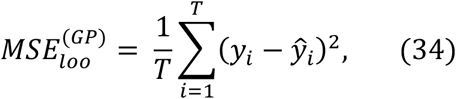

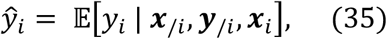

where 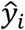 is the model prediction spike count for rendition ‘i’, and ***x***_/*i*_ and ***y***_/*i*_ are the song features and spike counts of all renditions excluding the i^th^ rendition. We then compared the GP model to a model with constant mean firing rate equal to the mean spike count over all renditions excluding the i^th^ rendition:

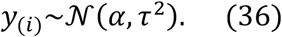

The predictive mean-squared error of this model is:

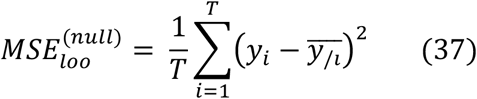

where,

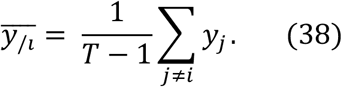

The cross-validated *r*^2^ value is:

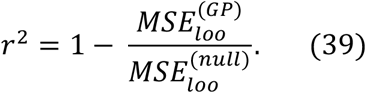

An *r*^2^ > 0 indicates that the model predicts new observations better than simply using the mean. The maximum theoretical value the *r*^2^ can take is one—this indicates perfect model prediction and, in practice, is never achieved. We use the *r*^2^ value as our measure of model performance.

#### Bootstrapping to assess population-level significance

Assessing the significance of the model predictions must be done on a population level for this type of analysis. We generated model fits to hundreds of song segment-spike count pairs for each syllable. Simply by chance, a portion of these fits would generate a predictive *r*^2^ > 0 value.

Furthermore, spike-song pairs are correlated, not only because of overlapping spike counts and song segment windows but also because of possible underlying correlations in the song and spike fluctuations across the song. To address this, we randomized the relationship between entire spike trains and song renditions and then re-performed our model fits on the randomized, spike count-song segment pairs across all time steps. By leaving the temporal structure of the song and spiking activity intact and only randomizing the relationship between them, we built a randomized population of fits for each cell-syllable pair, which retained the unknown, underlying temporal structure possibly present in the spike trains and song (Tusher et al., 2001). We repeated this procedure 100 times for the VTA_error_ cell population and 100 times for the VTA_other_ cell population to build a distribution of coherently randomized cell sets.

From this distribution of randomized cell sets, we computed single-tailed p-values assessments of the *r*^2^ values of the individual spike count-song segment model fits as well as on population measures of significance in the VTA_error_ and VTA_other_ cell populations independently. We assessed four population measures:

1. *The frequency of the predictive signal across the whole cell population.* An *r*^2^ > 0 indicates the model predicts the data better than an estimate based solely on the mean spike count across renditions, and we call this a ‘predictive signal’. We therefore assessed the significance of the total number of *r*^2^ > 0 song segment-spike count fits within the PPE latency window for both the VTA_error_ (p < 0.01) and VTA_other_ (p < 0.01) syllable-cell pairs respectively with a single-tailed p-value test.
2. *The spread of the predictive signal across the cell population.* We asked whether a small number of cells were accounting for the majority of the positive r^2^ values or if the signal appeared across multiple cells and syllables in the population. To answer this, we first labeled each cell-syllable pair as ‘significant’ if the number of positive r^2^ values within the PPE latency window (0-150 ms) had a single-tailed p < 0.05. We then calculated the single-tailed p-value for the number of significant cell-syllable pairs across the entire cell population (VTA_error_ population: one-sided z-test: p < 0.01; VTA_other_ population: one-sided z-test: p < 0.01).
3. *The significance of the magnitude of the peak in signal occurrence within the PPE latency window across the cell population.* We computed latency distributions as the latencies of the full set of spike-count song feature pairs that resulted in GP model fits with r^2^ > 0. We compared the variance of the latency distributions of the randomized populations to the variance of the peak we found in the actual data. We computed the single-tailed p-value for the maximum fluctuation of a latency distribution at any point in the latency domain. In this way we tested the significance not only of finding a peak in the data at the PPE window but of finding a peak of that size anywhere in the latency distribution. For example, VTA_error_ population latency distribution had a peak within the expected PPE latency region (Figure 3C). This peak was 3.84 standard deviations from the mean. The variance in relation to the maximum variance in randomized latency distributions was significant (one-sided z-test: p < 0.01). The time-bin of this latency distribution was 25 ms. VTA_other_ population latency distribution had a peak is 2.30 standard deviations from the mean (Figure 3D). This variance was not significant (one-sided z-test: p = 0.21). The time-bin of this latency distribution was 25 ms.
4. *The significance of the shapes of tuning curves we find via our GLM parameterization technique.* We computed the single-tailed p-value of the fraction of single-peaked tuning curves in the real population relative to the randomized populations (Figure 4E legend and main text).

Note that the goal of this significance strategy allows us to assess the VTA_error_ cell activity as a population, not the significance of particular song segment-spike count pairs. More data are needed for this level of specificity.

---

**Figure S1.**
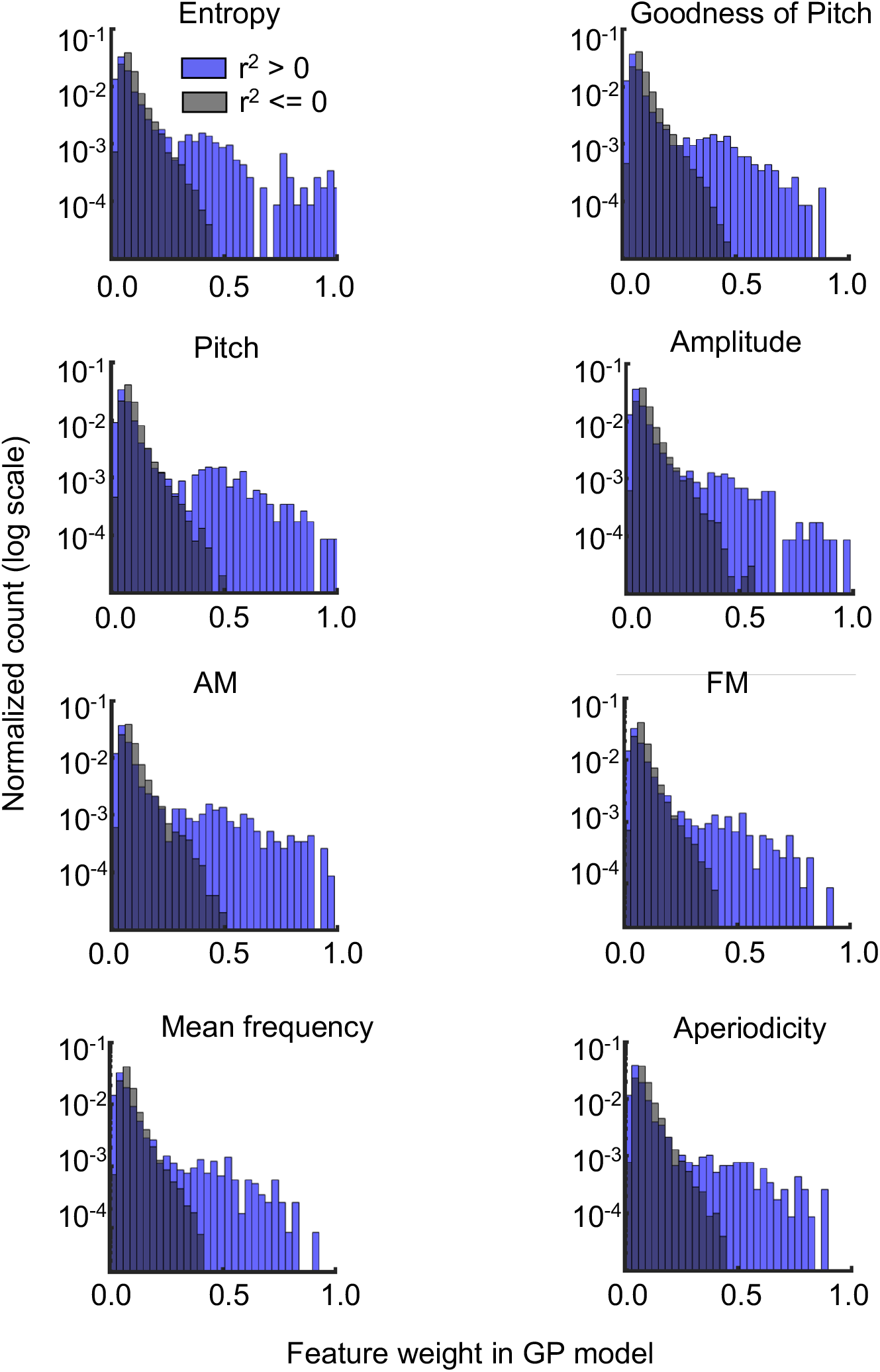
Related to Figure 2. Spiking Responses to Song Fluctuations are Captured by a Subset of Available Features, which Varies Across Context. Distributions of individual song feature weights in the full GP model for r^2^>0 and r^2^<=0 populations. In fits with r^2^>0 (predictive), the distribution was bimodal with an additional peak at around 0.5, implying that in predictive fits fewer numbers of features captured most of the information. In overfit models (r^2^<=0) all features contributed more equally to the poor estimate. Also note that the distributions across features were quite similar; no one feature captured significantly more of the song variations.

**Figure S2.**
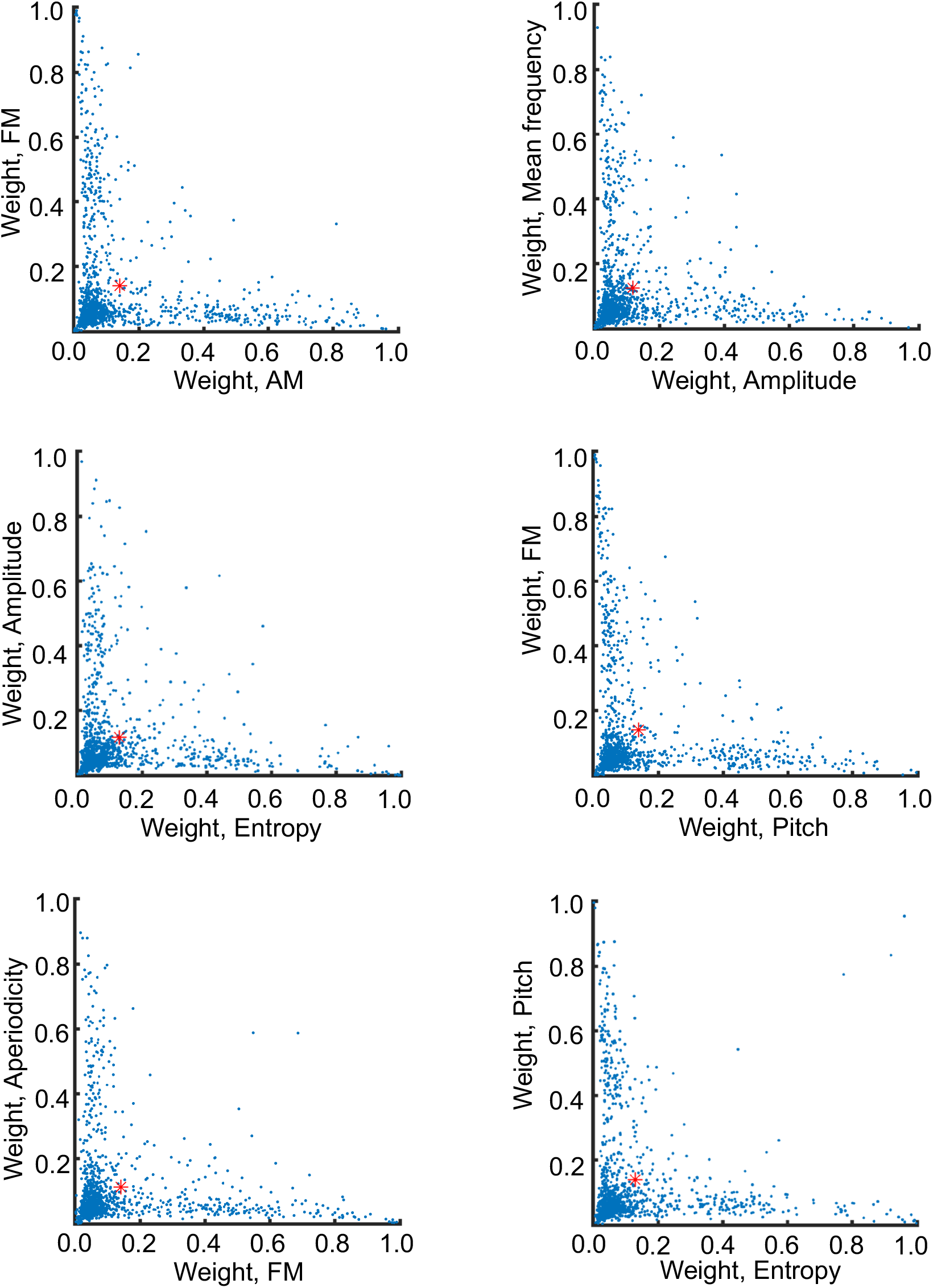
Related to Figure 2. Feature Contributions to Full GP Model Show Pairwise Correlations at Low, But Not High Weights. Scatterplots of relationships of feature contributions to the full GP model for individual fits with r^2^ > 0 within the PPE window. All plots: each point represents one model fit with an r^2^ > 0. Axes are the total feature weight in full the GP model of indicated feature; red dot is mean value. Pairs of features were selected to be representative of the full model set. All pairs showed comparable correlations at low weights.

**Figure S3.**
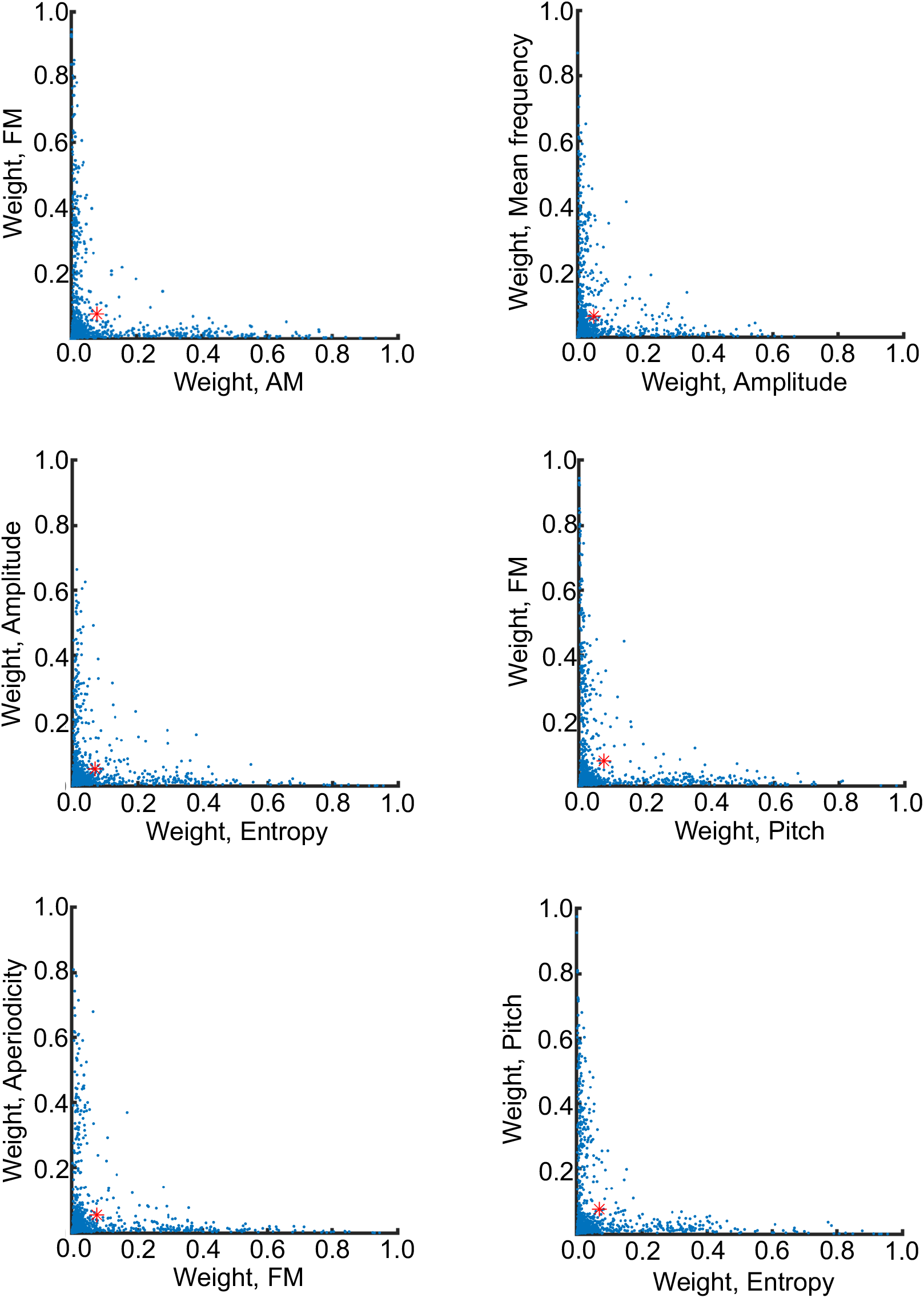
Related to Figure 2. 1^st^ Order Feature Contributions are Not Correlated. Scatterplots of Relationships of only the 1^st^ order feature contributions to the full GP model for individual fits with r^2^ > 0 within the PPE window. All plots: each point represents one model fit with an r^2^ > 0. Axes are the 1^st^ order feature weights in full GP model; red dot is mean value. Pairs of features were selected to be representative of the full model set. These data showed that correlations between features are mainly higher order coupling effects.

**Figure S4.**
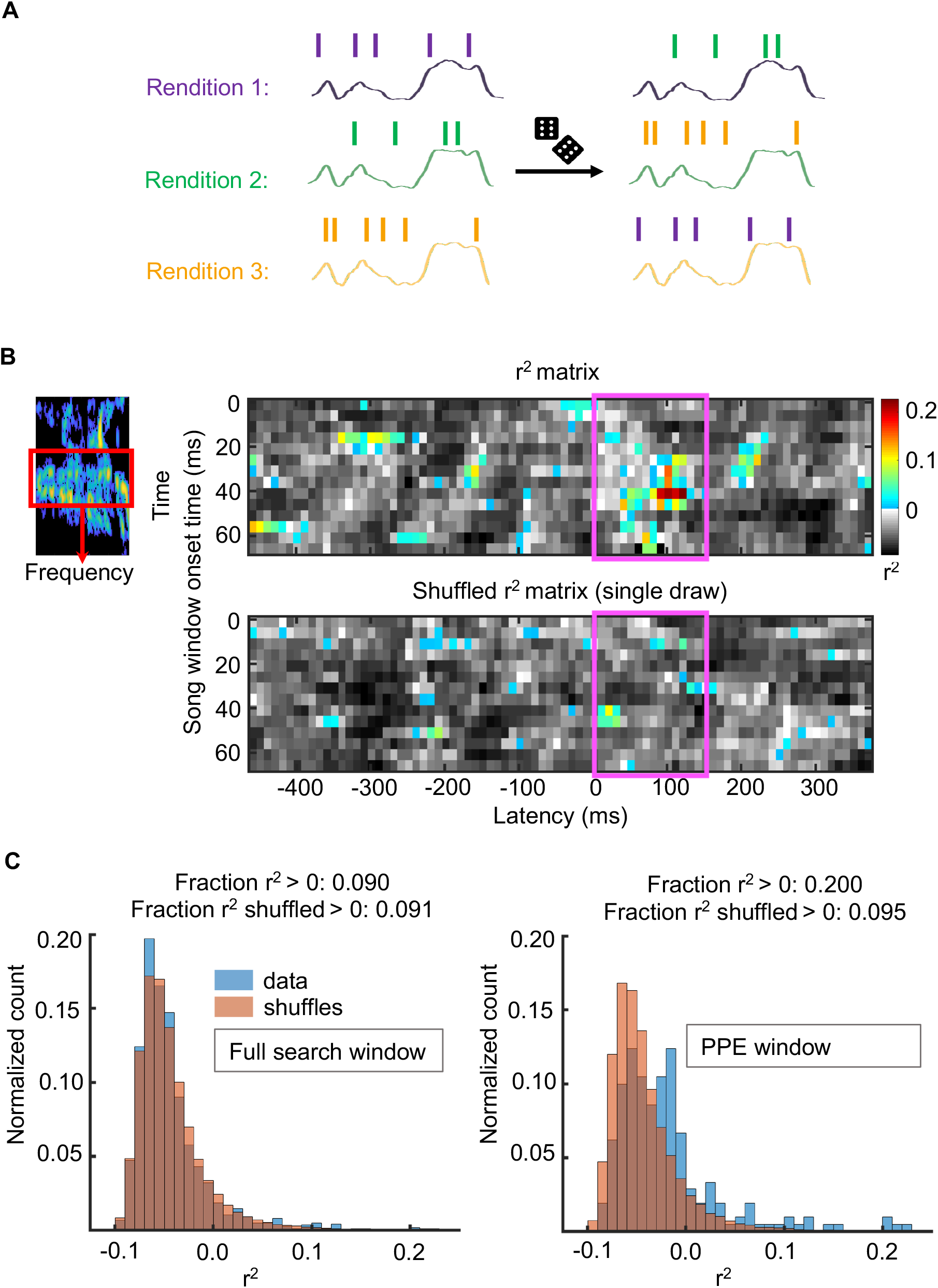
Related to Figure 3. Coherent Shuffling of Entire Spike Trains Retains Underlying Correlation Structure and Permits a Population-level Significance Assessment. (A) Schematic of spike train shuffling. Spike trains were randomized relative to the associated song. The randomized song-spike relationship was re-fit retaining possible underlying correlations in the spike train and song fluctuations. (B) Panel replicated from Figure 3A. (C) r^2^ distributions for randomized and actual data. Left: r^2^ distributions for the shuffled and real data compared across all latencies. The real and shuffled distributions appeared quite similar. The number of r^2^ > 0 in the real data was not significantly different from what would be expected by chance. Right: the distribution of r^2^ values that fall within the PPE latency window compared to the randomized distribution from within this same latency range. This distribution was shifted away from the randomized distribution, with more, larger r^2^ > 0. This population had a significantly greater number of r^2^ > 0 than expected by chance (one tailed z-test, p-value = 0.02).

